# Expression, purification and crystal structure determination of a ferredoxin reductase from the actinobacterium *Thermobifida fusca*

**DOI:** 10.1101/2020.03.12.988360

**Authors:** Jhon Alexander Rodriguez Buitrago, Thomas Klünemann, Wulf Blankenfeldt, Anett Schallmey

## Abstract

Ferredoxin reductase FdR9 from *Thermobifida fusca*, a member of the oxygenase-coupled NADH-dependent ferredoxin reductase (FNR) family, catalyzes electron transfer from NADH to its physiological electron acceptor ferredoxin. It forms part of a three-component cytochrome P450 monooxygenase system in *T. fusca*. Here, FdR9 was overexpressed and purified and its crystal structure was determined at 1.8 Å resolution. The overall structure of FdR9 is similar to other members of the FNR family and is composed of an FAD-binding domain, an NAD-binding domain and a C-terminal domain. Activity measurements with FdR9 confirmed a strong preference for NADH as the cofactor. Comparison of the FAD- and NAD-binding domains of FdR9 with other ferredoxin reductases revealed the presence of conserved sequence motifs in the FAD-binding domain as well as several highly conserved residues involved in FAD and NAD cofactor binding. Moreover, the NAD-binding site of FdR9 contains a modified Rossmann fold motif, GxSxxS, instead of the classical GxGxxG motif.

## 1. Introduction

Ferredoxin reductases (FdR) are essential components of the electron-transfer chains of three-component cytochrome P450 monooxygenases (Hannemann *et al.*, 2007). FdR9 from the actinobacterium *Thermobifida fusca* belongs to the oxygenase-coupled NADH-dependent ferredoxin reductase family of FAD-dependent electron-transfer enzymes [ferredoxin-NAD(P)H reductases; (EC 1.18.1.3)] (Vorphal et al., 2017). It mediates the transfer of two electrons from NADH to an iron-sulphur-cluster-containing ferredoxin via two successive one-electron transfer steps (Medina & Gómez-Moreno, 2004). Electron transfer between the ferredoxin reductase and the ferredoxin requires formation of a ternary NADH-FdR-Fdx complex (Deng et al., 1999; Kuznetsov et al., 2005). *T. fusca* is a moderately thermophilic soil bacterium (Bachmann & McCarthy, 1991) and a rich source of thermostable enzymes for application in biocatalysis (Wilson, 2004). In the genome of *T. fusca* (Lykidis et al., 2007), the gene coding for FdR9 (*Tfu_1273*) lies adjacent to the genes encoding CYP222A1 (*Tfu_1274*) and a ferredoxin named Fdx8 (*Tfu_1275*). Based on this genomic placement, FdR9, Fdx8 and CYP222A1 are believed to form a three-component cytochrome P450 monooxygenase system. Here, we report the heterologous production, purification, crystallization and structure determination of FdR9, one of the physiological protein partners of this three component system. The aim of this study was to provide a structural basis for further investigations of the intermolecular electron transfer processes between FdR 9 and Fdx8 as well as the associated protein-protein interactions.

## 2. Materials and methods

### 2.1. Macromolecule production

The gene encoding FdR9 (*Tfu_1273*) was provided by Prof. Vlada Urlacher (Institute of Biochemistry II, Heinrich-Heine University, Düsseldorf, Germany) in the plasmid pET22b(+). The gene was subcloned from pET22b(+) into vector pET28a(+) using restriction enzymes NdeI and EcoRI for N-terminal His_6_-tag fusion of the resulting protein and the presence of a thrombin recognition site between His-tag and the FdR9 sequence. FdR9 was overexpressed in *Escherichia coli* C43 (DE3) (Miroux & Walker, 1996; Dumon-Seignovert et al., 2004) cells transformed with the *Tfu_1273*-containing pET28a(+) vector. For functional expression, bacteria were grown in 1 L TB medium supplemented with 50 mg L^-1^ kanamycin at 37 °C. At OD_600_ = 0.8, expression was induced by the addition of IPTG to a final concentration of 0.5 mM. After expression for 16 hours at 240 rpm and 37 °C, the culture was harvested by centrifugation. (15 min at 4400 g, 4 °C) and resuspended in lysis buffer (20 mM Tris pH 8.0, 300 mM NaCl), always in the presence of protease inhibitors (1 tablet *Complete EDTA-free* per 50 mL; Sigma-Aldrich). After sonication (9 cycles of 20 s at an amplitude of 60% followed by 10 s pause, on ice using a Vibra-Cell™ VCX130, Sonics & Materials, USA) for cell disruption and subsequent centrifugation (15 min, 10000 g, 4 °C) to remove cell debris, FdR9 was found in the soluble fraction. The His_6_-tag-containing FdR9 was purified by affinity chromatography on an Äkta prime FPLC system (GE Healthcare, Freiburg, Germany), using a 5 mL HisTrap column (GE Healthcare). The bound protein was eluted using a linear imidazole gradient (0 - 0.5 M) in 5 column volumes (CV). Selected FdR9-containing fractions were combined for incubation with the Thrombin CleanCleave™ Kit (Sigma Aldrich) and afterwards dialyzed against 20 mM Tris pH 8.0, 40 mM NaCl, at 4 °C for 16 hours. Dialyzed samples were loaded again on a HisTrap column (GE Healthcare) to remove the cleaved His-tag. The FdR9-containing flow-through was loaded on a 5 mL HiTrap Q HP column (GE Healthcare) and elution was done using a linear NaCl gradient (0.04 - 1 M) in 5 CV. Selected FdR9-containing fractions were concentrated by ultrafiltration using a 30 kDa cut-off membrane (Amicon ultra-15, Merck) and further purified by gel filtration on a Superdex 75 26/60 column (GE Healthcare) using 20 mM Tris pH 8.0, 300 mM NaCl, 1 mM DTT. Purified FdR9 was obtained with a yield of 60 mg L^-1^ culture and displayed a yellow colour indicative of the flavin cofactor. A high degree of purity was confirmed for FdR9 by observation of a single band on a 12% SDS-PAGE gel.

### 2.2. Crystallization

Initial crystallization trials were carried out at room temperature in 96-well INTELLI-Plates (Art Robbins Instruments, Sunnyvale, CA, USA) with Index Screen (D’arcy et al., 2003) using the sitting drop vapor-diffusion method. The droplet was initially formed by 200 nL of a solution containing 42 mg mL^-1^ FdR9 in 20 mM Tris pH 8.0, 300 mM NaCl, mixed with 200 nL of reservoir solution using a pipetting robot (Honeybee 963, Genomic Solutions, Huntingdon, U.K) and then equilibrated against 60 µL of reservoir solution. Crystals of FdR9 were obtained in the condition Index C6 containing 1.5 M ammonium sulfate, 0.1 M NaCl and 0.1 M BIS-Tris at pH 6.5. The crystals were yellow-colored as is typical for native FNR crystals due to the presence of the flavin cofactor (Morales et al., 2000) (Fig. 1). Trials to optimize crystal quality by variation of precipitant concentration or pH did not result in better diffracting crystals.

**Figure 1.**
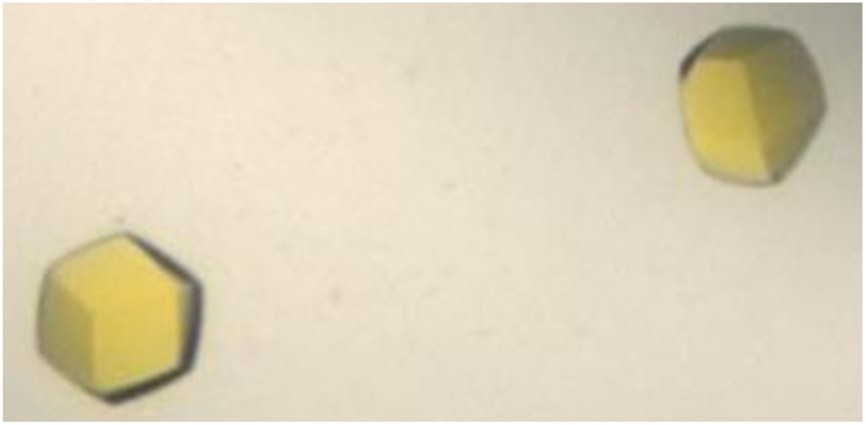
Crystals of ferredoxin reductase FdR9 from *Thermobifida fusca*, with size dimensions (50 × 50 × 50 µm) obtained from the Index C6 condition.

### 2.3. Data collection and processing

FdR9 crystals were harvested using a nylon loop (Hampton Research) and soaked in reservoir solution containing 20% (v/v) 2,3-(*R,R*)-butandiol prior to flash freezing in liquid nitrogen. 3600 images were collected using the oscillation method with a range of 0.1^°^ per image on a Dectris Pilatus 6M-F detector using single wavelength synchrotron radiation at the beamline X11 at PETRAIII of the Deutsches Elektronen-Synchrotron (DESY, Hamburg, Germany). Reflection image processing was performed using DIALS (Winter et al., 2018), AIMLESS (Evans & Murshudov, 2013) of the CCP4 suite (Collaborative, 1994). Data collection and processing statistics are summarized in Table 1.

**Table 1.**
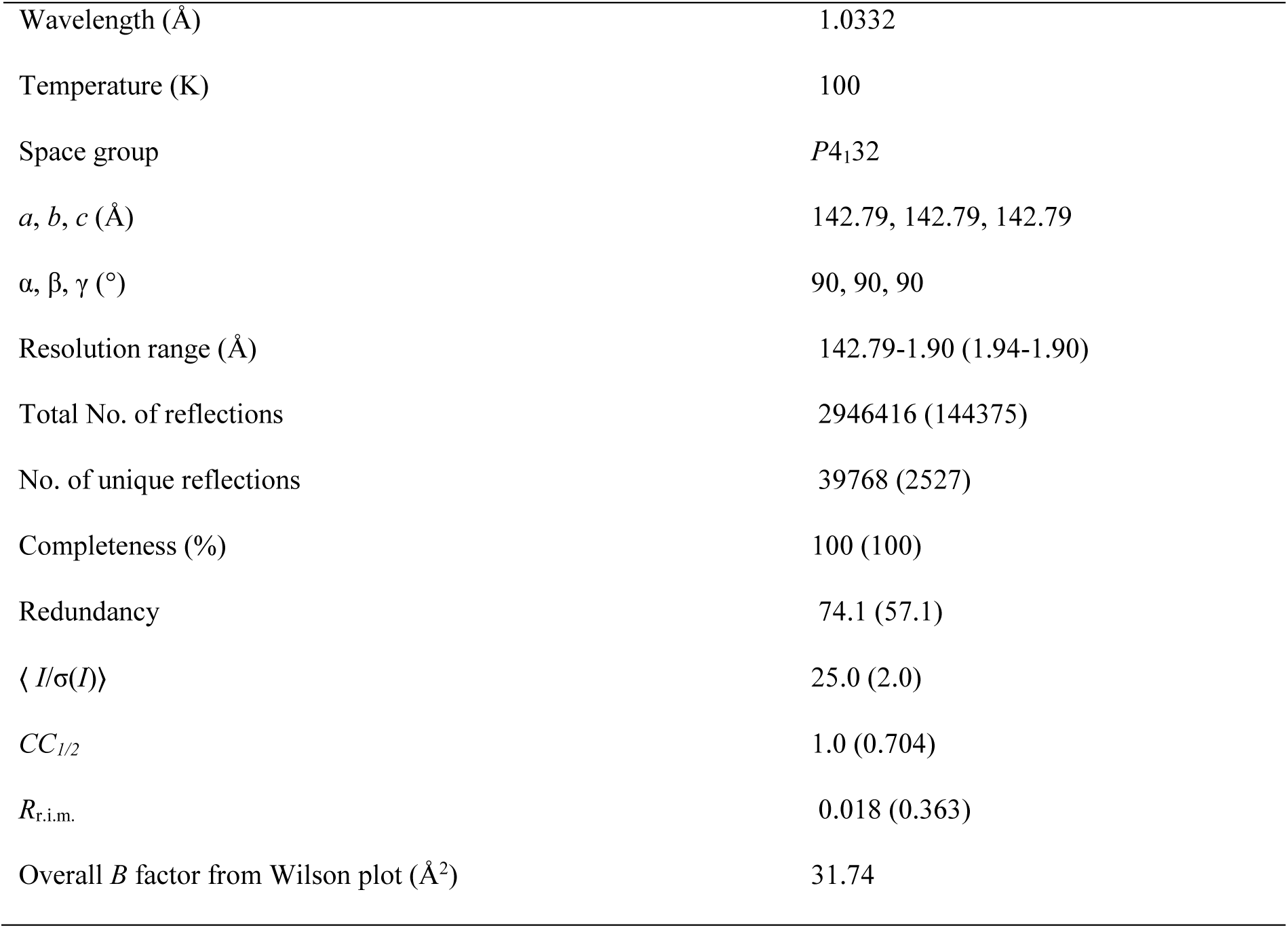
Data collection and processing Values for the outer shell are given in parentheses.

|  |  |
| --- | --- |
| Wavelength (Å) | 1.0332 |
| Temperature (K) | 100 |
| Space group | *P*4<sub>1</sub>32 |
| *a*, *b*, *c* (Å) | 142.79, 142.79, 142.79 |
| $\alpha$, $\beta$, $\gamma$ (°) | 90, 90, 90 |
| Resolution range (Å) | 142.79–1.90 (1.94–1.90) |
| Total No. of reflections | 2946416 (144375) |
| No. of unique reflections | 39768 (2527) |
| Completeness (%) | 100 (100) |
| Redundancy | 74.1 (57.1) |
| $\langle I/\sigma(I) \rangle$ | 25.0 (2.0) |
| *CC*<sub>1/2</sub> | 1.0 (0.704) |
| *R*<sub>r.i.m.</sub> | 0.018 (0.363) |
| Overall *B* factor from Wilson plot (Å<sup>2</sup>) | 31.74 |

### 2.4. Structure solution and refinement

Initial phases were obtained by molecular replacement using Mr.BUMP (Keegan & Winn, 2008) executing PHASER (McCoy *et al.*, 2007) and using the atomic coordinates of putidaredoxin reducatse (PDB: 1q1w) (Sevrioukova et al., 2004) as search model. Refinement was performed by alternating rounds of REFMAC5 (Murshudov et al. 2011) and manual adjustments in COOT (Emsley et al., 2010). Last refinement steps were performed with phenix.refine (Afonine et al., 2005) including TLS refinement and the addition of riding hydrogens. FdR9 diffraction data and coordinates were deposited in the Protein Data Bank (Berman et al., 2002) (PDB entry: 6TUK). Representations of the structures were generated with PyMOL Molecular Graphics System version 2.1.1 (Schrödinger, LLC, New York, NY, USA). Refinement statistics are listed in Table 2.

**Table 2.**
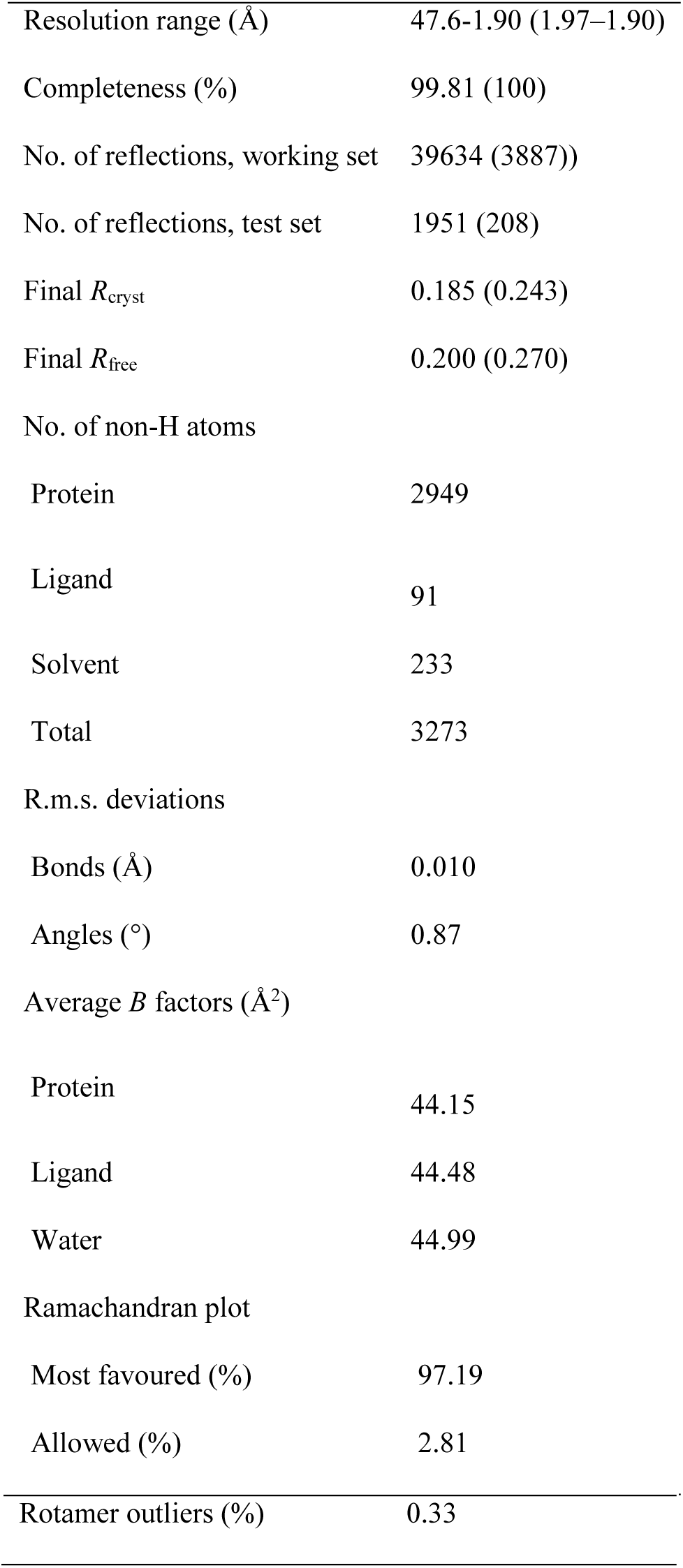
Structure solution and refinement Values for the outer shell are given in parentheses.

|  |  |
| --- | --- |
| Resolution range (Å) | 47.6–1.90 (1.97–1.90) |
| Completeness (%) | 99.81 (100) |
| No. of reflections, working set | 39634 (3887) |
| No. of reflections, test set | 1951 (208) |
| Final $R_{\text{cryst}}$ | 0.185 (0.243) |
| Final $R_{\text{free}}$ | 0.200 (0.270) |
| No. of non-H atoms |  |
| Protein | 2949 |
| Ligand | 91 |
| Solvent | 233 |
| Total | 3273 |
| R.m.s. deviations |  |
| Bonds (Å) | 0.010 |
| Angles (°) | 0.87 |
| Average $B$ factors (Å<sup>2</sup>) | |
| Protein | 44.15 |
| Ligand | 44.48 |
| Water | 44.99 |
| Ramachandran plot |  |
| Most favoured (%) | 97.19 |
| Allowed (%) | 2.81 |
| Rotamer outliers (%) | 0.33 |

### 2.5. Reductase activity assay

The activity of FdR9 with NADH and NADPH as cofactors was determined spectrophotometrically by measuring the decrease in ferricyanide concentration at 420 nm (ε_420_ = 1.02 mM^-1^ cm^-1^) (Roome et al., 1983). Each 1 mL reaction contained 0.5 mM K_3_Fe(CN)_6_, an appropriate amount of FdR9 (3.7 µg for the reaction with NADH, 37 µg for the reaction with NADPH) and 0.5 mM NADH or NADPH in 50 mM potassium phosphate buffer, pH 7.4. Measurements were performed at ambient temperature on a Cary 60 instrument (Agilent, Heilbronn, Germany).

## 3. Results and discussion

### 3.1. Structural overview of FdR9

FdR9 crystallized in space group P4_1_32 with one monomer in the asymmetric unit. Initial phases were obtained by molecular replacement using the structure of putidaredoxin reductase (PDB code 1q1w; 26% amino-acid sequence identity) as the search model. The resulting electron density map allowed identification of the FAD molecule bound to the reductase. The final model of FdR9, refined to R_cryst_ of 18.5% and a R_free_ of 20.0%, displays very good geometry with no residues located in disallowed regions of the Ramachandran plot. The structure of FdR9 contains nine α-helices and 25 β-strands which form three distinct domains (Fig. 2): an FAD-binding domain (residues 1–106 and 223–308), an NAD-binding domain (residues 107–222), and a C-terminal domain (residues 309–393). Structural comparison of FdR9 with other proteins using the DALI server (Holm & Sander, 1995) indicated that it is very similar to ferredoxin reductases from *Rhodopseudomonas palustris* (PuR), *Novosphingobium aromaticivorans* (ArR), *Pseudomonas* sp. (BphA4) and *Pseudomonas putida* (PdR) (Table 3), all sharing the three-domain architecture as shown in Figure 2. Root-mean-square deviations (r.m.s.d.) for Cα atoms of these protein structures to the structure of FdR9 ranged between 1.9 to 2.2 ß. PuR, ArR and PdR are part of three-component cytochrome P450 monooxygenase systems like FdR9, whereas BphA4 forms part of the three-component biphenyl dioxygenase system present in *Pseudomonas* sp. (Table 3).

**Table 3.** DALI search against the PDB using the FdR9 structure

| PDB id | Organism | Terminal oxygenase | Alignment length | Dali Z-score | Reference |
| --- | --- | --- | --- | --- | --- |
| 3FG2<br>(PuR) | *Rhodopseudomonas palustris* | CYP199A2 | 380 | 46.8 | (Xu *et al.*, 2009) |
| 3LXD<br>(ArR) | *Novosphingobium aromaticivorans* | CYP101D1 | 383 | 47.5 | (Yang *et al.*, 2010) |
| 2GQW<br>(BphA4) | *Pseudomonas* sp. | Biphenyl dioxygenase<br>BphA1A2 | 382 | 47.2 | (Senda *et al.*, 2007) |
| 1Q1W<br>(PdR) | *Pseudomonas putida* | P450cam | 386 | 46.6 | (Sevrioukova *et al.*, 2004) |

**Figure 2.**
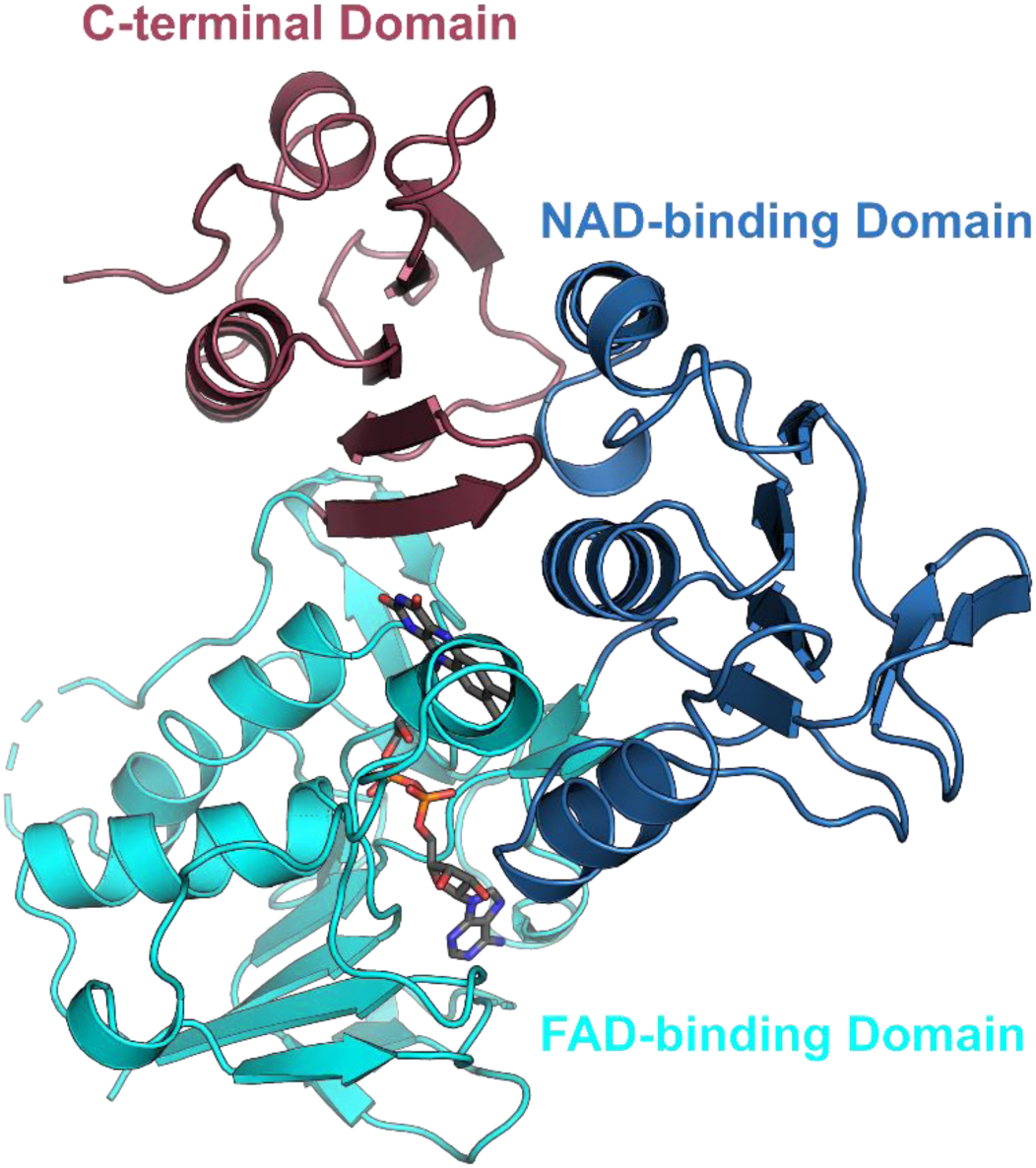
Overall structure of FdR9. The FAD-binding, NAD-binding, and C-terminal domains are colored in cyan, light blue, and light pink, respectively.

### 3.2. FAD-binding domain

The FAD-binding domain of FdR9 adopts a typical α/β-fold consisting of two antiparallel β-sheets and one parallel β-sheet surrounded by four α-helices, as shown in Figure 2. Sequence alignment analysis of the FAD-binding domain reveals the presence of three highly conserved motifs PYxRPPLSK, TSxPx(3)AxG and RxEx(4)A (Fig 3). These motifs contain most of the amino acids that interact with the FAD cofactor. In FdR9, the FAD cofactor interacts with the protein through a network of hydrogen bonds on the FAD-binding domain involving amino acid residues Leu11, Ala12, Glu35, Arg42, Lys47, Ala76, Arg119, Asp260 and Trp278 (Fig. 4), of which several are conserved among the ferredoxin reductases PuR, ArR, PdR and BphA4 (Fig. 3). The pyrophosphate moiety of FAD is stabilized by hydrogen bonds between main chain amide groups of Ala12 and Leu11 with O_1_P, the side chain NH_2_ group of Arg119 with O_2_A as well as the Asp260 side chain hydrogen bonding with the O_2_P atom and O_3_′ of the riboflavin part. Moreover, Arg42 plays an important role in stabilizing the FAD cofactor through the formation of electrostatic interactions and hydrogen bonds with the O_1_A atom of the pyrophosphate moiety and the O_3_B atom of the adenosine ribose (Fig. 4). In PdR, this arginine is replaced by Leu45, while PuR, ArR and BphA4 also carry an arginine at the respective position (Fig. 3 and 4). Ala76 of FdR9 is involved in the stabilization of the adenine through hydrogen bonding of the backbone carbonyl oxygen with the N_6_A atom. The Oε of Glu35 forms hydrogen bridges with atoms O_2_B and O_3_B of the adenosine ribose. This glutamate is also conserved in PuR, ArR and BphA4, whereas an alanine is present at the corresponding position in PdR (Fig. 3 and 4). Residue Trp278 stabilizes the isoalloxazine ring of the FAD cofactor by hydrogen bonding between its backbone NH and O_2_ of the isoalloxazine ring. Lys47 in FdR9 forms a salt bridge with Glu148 and hydrogen bonds with the carbonyl oxygen of Pro43 as well as with the O_4_ and N_5_ atoms of the isoalloxazine ring.

**Figure 3.**
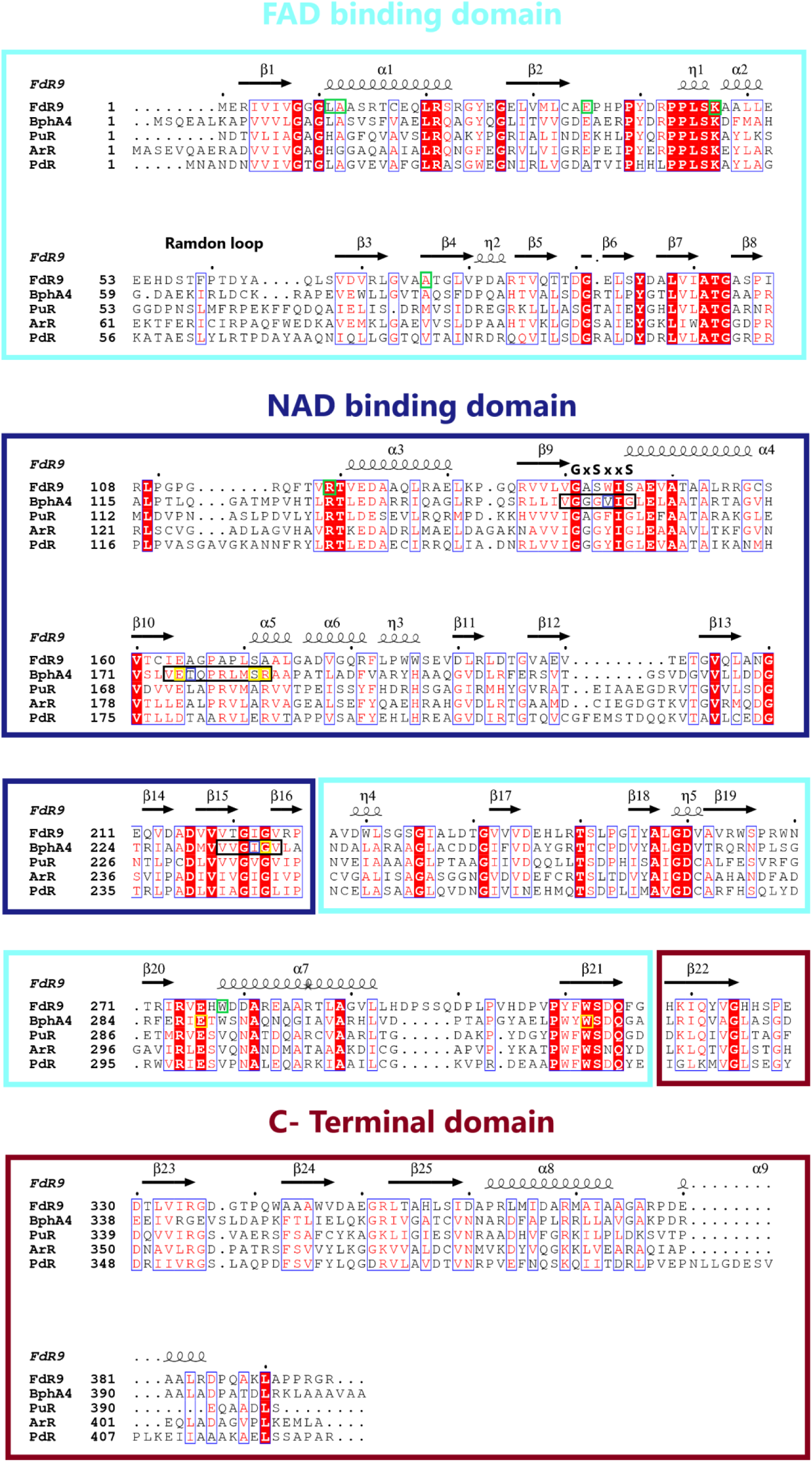
Multiple sequence alignment of FdR9 and homologous ferredoxin reductases, obtained in the DALI search using th**e** FdR9 structure as template. Helices are represented by co**i**ls and β-sheets are shown as arrows. Columns with residues that are more than 70% similar according to physico-chemical properties (threshold set to 0.7) are framed in blue, amino acid residues with 100% identity are highlighted by red background. FdR9 residues involved in hydrogen bond interaction with the FAD molecule are framed in green boxes. The three loops of BphA4 interacting with the NAD molecule are framed in black boxes, and the residues of BphA4 interacting with the NAD molecule by hydrogen bonding and hydrophobic interactions are highlighted by yellow boxes. The PAM250 matrix was used for sequence alignment. The figure was rendered by ESPript 3.0 (Robert & Gouet, 2014).

**Figure 4.**
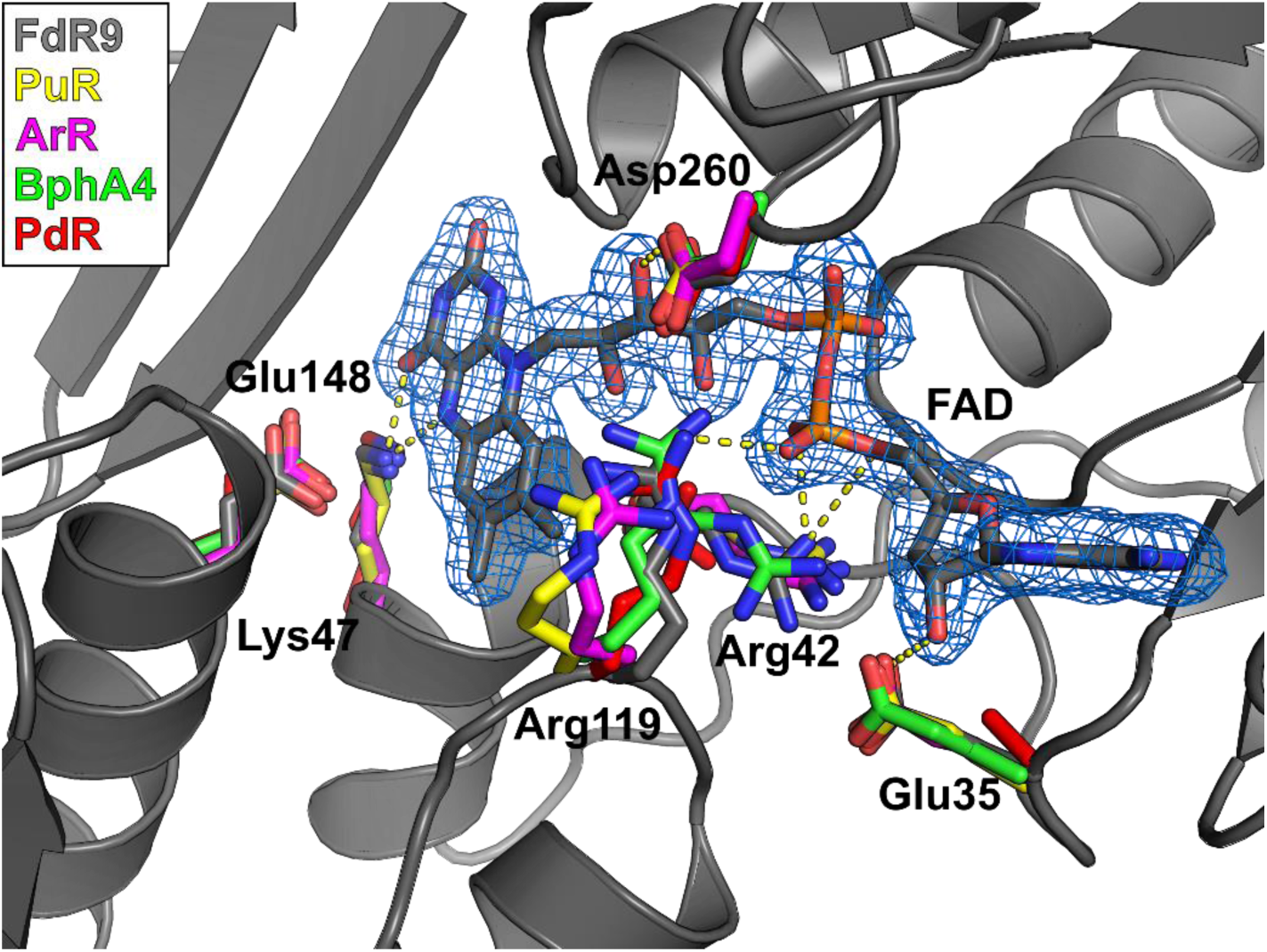
Superposition of the FAD-binding site of FdR9 (PDB: 6TUK, gray color) with structures of other ferredoxin reductases [PDB IDs: 3FG2 (PuR), 3LXD (ArR), 2GQW (BphA4) and 1Q1W (PdR)]. Residues forming hydrogen bond interactions with the FAD cofactor are shown as sticks. Hydrogen bonds are indicated by dotted lines.

Despite the fact that the overall fold of the FAD domain in FdR9 is highly similar to structures of PuR, ArR, PdR and BphA4 with several conserved amino acids interacting with the FAD molecule, differences can be observed as well. The first difference concerns the presence or absence of secondary structure in the random loop region (residues 51-63 in FdR9) of the five ferredoxin reductases. The short and longer helices found in this region in PdR, PuR, ArR and BphA4 correspond to a purely random loop in FdR9 (Fig. 5). It has previously been proposed that the absence of helix secondary structure in this region could influence the binding of FAD (Xu *et al.*, 2009). Another difference is the insertion of five amino acids (residues 295-300) in a surface loop in FdR9, while the loop is shorter in the other four reductases (Fig. 3). The electron density for residues 298-300 in this extended loop in FdR9 is not well resolved indicating high flexibility.

**Figure 5.**
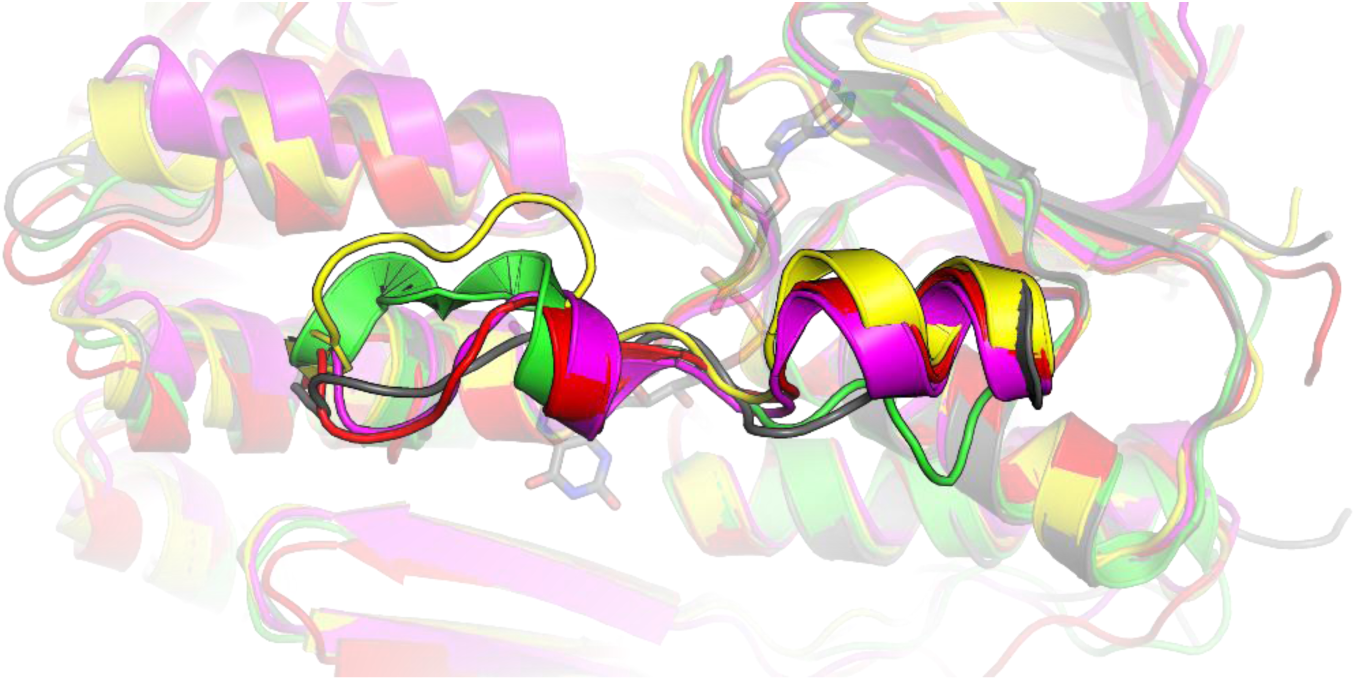
Superposition of the structural model of FdR9 (PDB: 6TUK, in gray) with structures of other ferredoxin reductases [PDB IDs: 3FG2 (PuR) in yellow, 3LXD (ArR) in magenta, 2GQW (BphA4) in green and 1Q1W (PdR) in red] to highlight differences in secondary structure in the random loop region (residues 51-63 in FdR9).

### 3.3. NAD-binding domain

In our FdR9 crystallization experiments we did not attempt to co-crystallize FdR9 with nicotinamide cofactor and hence, the structure of FdR9 presented here does not contain NAD. Nevertheless, sequence and structure analysis of the NAD-binding domain of FdR9 indicates the presence of a canonical Rossmann fold similar to previously reported complexes, where the NAD molecule was shown to interact with three loops [corresponding to residues 140-146 (first loop), 164-172 (second loop) and 219-224 (third loop) in FdR9; Fig. 3] (Senda et al., 2000). In PuR, ArR, PdR and BphA4, the first loop between β9 and α4 contains the typical GXGXXG motif indicative of nicotinamide cofactor binding, which is modified to **G**A**S**WI**S** in FdR9 with the last two glycines of the motif replaced by serine (Fig. 3). This change in the motif does not affect the folding of the respective loop as can be seen in the structural comparison between FdR9 and BphA4 (Fig 6). Previous studies proposed that the sequence motif GXGXXG was indicative of NAD specificity, whereas the motif GXGXXA is found in NADP-binding enzymes, although exceptions are known (Carugo & Argos, 1997; Hanukoglu, 2017), as would be the case for FdR9. In our studies, FdR9 was found to display a clear preference for NADH as the cofactor. Based on a ferricyanide reduction assay (Roome et al., 1983), the specific activity of FdR9 with NADH was determined as 104 U mg^-1^, whereas the specific activity with NADPH was only 1.2 U mg^-1^. Hence, the nicotinamide cofactor-binding site of FdR9 was compared in more detail to that of BphA4 (PDB: 1F3P), which has been crystallized with NAD bound in the active site. The comparison shows that FdR9 shares a similar fold at the entrance of the NAD-binding channel and an accessible nicotinamide-binding site above the isoalloxazine ring of the flavin cofactor. In BphA4, NAD interacts with residues Val155, Ile156 and Glu159 in the first loop (corresponding residues in FdR9 are Trp144, Ile145 and Glu148) and residues Glu175, Thr176, Ser182 and Arg183 in the second loop (corresponding to residues Glu164, Ala165, Ser171 and Ala172 in FdR9). Glu175 of BphA4 forms hydrogen bonds with the O_2_B and O_3_B atoms of the adenosine ribose as well as with the amino acid Thr176. The negative charge of Glu175, which is not only conserved in FdR9 (Fig. 6), but also in PuR, ArR and PdR, has been proposed to enhance NAD cofactor specificity by repulsion of the phosphate of NADP (Hanukoglu, 2017). Ser182 in BphA4, which corresponds to Ser171 in FdR9, forms a hydrogen bond with the adenosine ribose O_3_B atom, too. Residues in the third loop interacting with the NAD molecule (Ile235 and Gly236 in BphA4) as well as Glu289 and Trp320 of BphA4 are significantly conserved among the five ferredoxin reductases compared in Figure 3. Interestingly, FdR9 carries a threonine at position 220 (third loop), while PuR, ArR, PdR and BphA4 possess a hydrophobic residue (either valine or alanine) at the corresponding position. Of the 11 amino acids directly interacting with the NAD molecule in BphA4 by hydrogen bonding, eight are conserved in FdR9 (Fig. 3), whereas both proteins share an overall amino acid sequence identity of only 30%. In contrast, other loop residues (especially in the first and second loop), which are not forming direct hydrogen bond interactions with the nicotinamide cofactor, display a lower degree of conservation.

**Figure 6.**
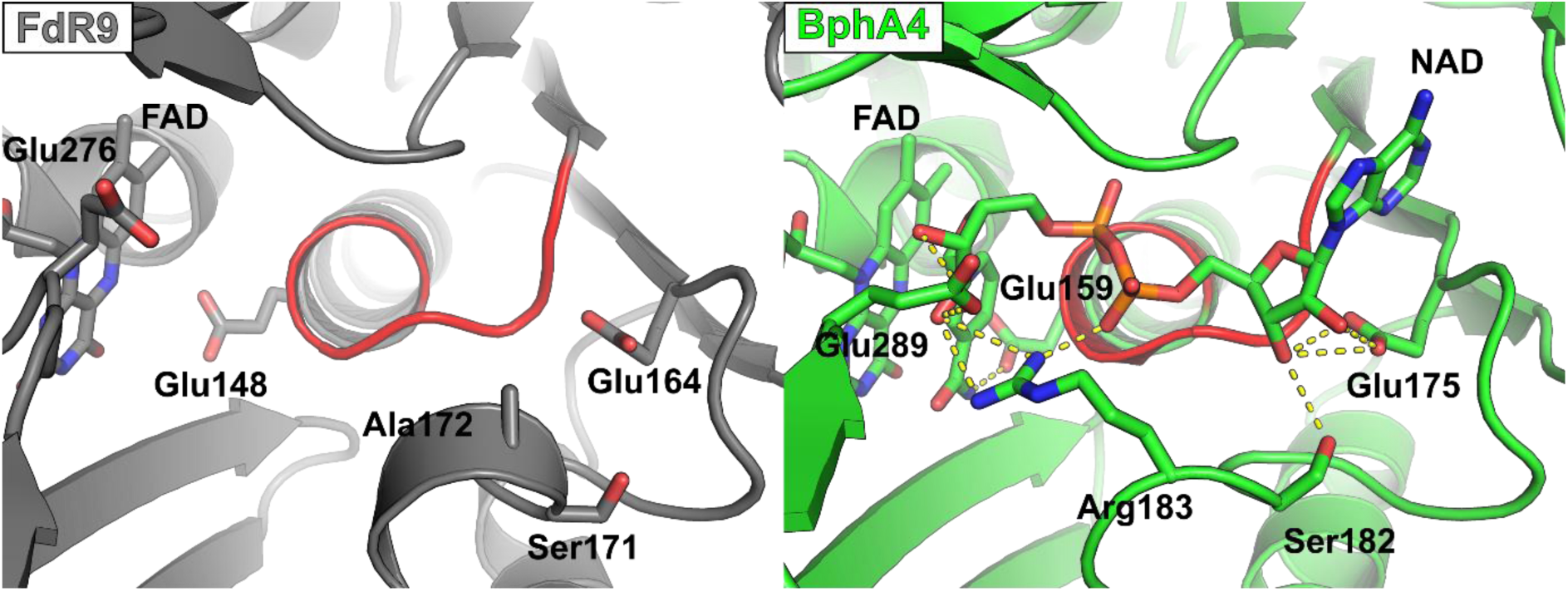
Structural comparison of the NAD-binding sites of FdR9 (PDB: 6TUK) and the reductase BphA4 (PDB: 1F3P) in complex with NAD^+^. The amino acid residues of BphA4 interacting with NAD^+^ and the corresponding residues in FdR9 are labeled. The loops containing motif GxGxxG of BphA4 and motif GxSxxS of FdR9 are shown in red color.

Apart from this, FdR9 exhibits two significantly shorter surface loops in the NAD-binding domain compared to the other four ferredoxin reductases. This regards the loops between β8 and α3 as well as β12 and β13 (Fig. 3). Especially the latter is 7, 7 and 9 amino acids longer in PuR, ArR and PdR, respectively. In PdR, this loop is involved crystallographic dimer formation, while PuR and ArR crystallize as monomers like FdR9 (Sevrioukova et al., 2004; Xu et al., 2009; Yang et al., 2010). It is unclear, however, if those differences in surface loop lengths also have a possible physiological impact.

## Acknowledgements

We thank the beamline staff at P11 at the PETRAIII synchrotron (Deutsches Elektronensynchrotron DESY, Hamburg, Germany) for granting us access to their facilities. This project was financially supported by the Deutsche Forschungsgemeinschaft (DFG, German Research Foundation) via the Research Training Group PROCOMPAS [GRK 2223].

